# GRASP: a Bayesian network structure learning method using adaptive sequential Monte Carlo

**DOI:** 10.1101/767327

**Authors:** Kaixian Yu, Zihan Cui, Xing Qiu, Jinfeng Zhang

## Abstract

Bayesian networks (BNs) provide a probabilistic, graphical framework for modeling high-dimensional joint distributions with complex dependence structures. BNs can be used to infer complex biological networks using heterogeneous data from different sources with missing values. Despite extensive studies in the past, network structure learning from data is still a challenging open question in BN research. In this study, we present a sequential Monte Carlo (SMC) based three-stage approach, GRowth-based Approach with Staged Pruning (GRASP). A double filtering strategy was first used for discovering the overall skeleton of the target BN. To search for the optimal network structures we designed an adaptive SMC (adSMC) algorithm to increase the diversity of sampled networks which were further improved by a new stage to reclaim edges missed in the skeleton discovery step. GRASP gave very satisfactory results when tested on benchmark networks. Finally, BN structure learning using multiple types of genomics data illustrates GRASP’s potential in discovering novel biological relationships in integrative genomic studies.

## 1. Introduction

A Bayesian network (BN) is a graphical representation of the joint probability distribution of a set of variables (called nodes in the graph). BNs have been widely used in various fields, such as computational biology (Friedman, Linial, Nachman and Pe’er 2000, Raval, Ghahramani and Wild 2002, Vignes, et al. 2011), document classification (Denoyer and Gallinari 2004), and decision support system (Kristensen and Rasmussen 2002). The availability of large volumes of genomics data have made it possible to learn the complex biological networks governing the interactions of various biomolecules. BNs are powerful tools for learning biological networks (Yu, Smith, Wang, Hartemink and Jarvis 2004) and it offers several advantages compared to other methods. First, BNs learn causal relationships, which will help researchers understand the regulatory relationships among different bio-entities; Second, BNs provide a probabilistic framework, which can be easily interpreted and integrated with other analysis; Third, BNs can easily integrate heterogeneous data of different types or from different sources; Fourth, BNs can conveniently deal with missing values in the data.

BN encodes conditional dependencies and independencies (CDIs) among variables into a directed acyclic graph (DAG). And this DAG is called the structure of a BN. When the structure of a BN is given, the parameters that quantify the conditional dependencies can be estimated from the observed data. If neither the parameters nor structures are given, they can be inferred from observed data. In this study, we will be focusing on the structure estimation (or learning) of a BN.

The technical difficulties of structure learning lie on the fact that the DAG space is of super-exponential cardinality and is quite rugged for most commonly used score functions. Estimating the global optimal structure given an observed dataset exactly is an NP-hard problem (Cooper 1990, Koller and Friedman 2009). There have been many inexact and heuristic methods proposed in the past two decades. The strategy of these methods can be classified mainly into three categories: constraint based, score based, and hybrid, which combines both constraint and score based approaches.

A constraint based method utilizes a suitable conditional dependency test to identify the conditional dependencies and independencies among all nodes (Campos 1998, de Campos and Huete 2000, Margaritis 2003, Tsamardinos, Aliferis and Statnikov 2003, Yaramakala and Margaritis 2005, Aliferis, Statnikov, Tsamardinos, Mani and Koutsoukos 2010). A major disadvantage of such a method is that a large number of tests have to be conducted; therefore, an appropriate method to adjust the *p*-values obtained from all the tests is desired, which reduces the statistical power of detecting conditional dependencies. The fact that not all the tests are mutually independent makes the *p*-values adjustment even more difficult. Another issue is that the goodness-of-fit of the obtained network has not been considered; therefore, the estimated BN may not fit the observed data well.

A score based method uses a score function to evaluate the structures of BNs on training data (Larrañaga, Poza, Yurramendi, Murga and Kuijpers 1996, Friedman, Nachman and Peér 1999, Gámez, Mateo and Puerta 2011). A searching algorithm is employed to search the best BN (with the optimal score) with respect to certain score function. Various Bayesian and non-Bayesian score functions have been proposed in the past. As exact search is not feasible, over the past two decades, various heuristic searching methods, such as Hill climbing, tabu search, and simulated annealing were proposed to search for the optimal BN structures. The problem with score based method is that the search space is often very large and complicated; therefore, the searching algorithm either will take too much time to find the optimum or be trapped in local optima. Many efforts have been made to overcome this challenging issue, such as searching using an ordered DAG space to reduce the search space (Teyssier and Koller 2012). In the ordered DAG space, the nodes are given an order such that edges will only be searched from higher order to lower order. The practical issue is that determining the order and finding the optimal structure is equally difficult. More recently, various penalty based methods were proposed to estimate the structures for Gaussian BN (GBN) (Fu and Zhou 2013, Huang, et al. 2013, Xiang and Kim 2013). These methods have been shown to be quite efficient for GBN structure learning and are able to handle structure learning and parameter estimation simultaneously; however, these methods are quite restrictive: the joint distributions must approximately follow a multivariate Gaussian distribution and dependencies among nodes are assumed to be linear.

Hybrid methods which combine a constraint method and a score based method were proposed to combine the advantages of both methods (Tsamardinos, Brown and Aliferis 2006). Such methods often contain two stages: first pruning the search space by constraint based methods, then searching using a score function over the much smaller pruned space. In the pruning stage, the goal is to identify the so-called skeleton of the network, which is the undirected graph of the target DAG. Later in the second stage, the direction of each edge will be determined by optimizing the score function. In a hybrid method, it is important that the first stage identifies as many true undirected edges as possible, since only the identified undirected edges will be considered in the second stage.

In this study, we developed a novel BN structure learning method, named GRASP (GRowth-based Approach with Staged Pruning). It is a three-stage method: in stage one, we used a double filtering method to discover a cover of the true skeleton. Unlike the traditional constraint methods, which try to obtain the true skeleton exactly, our method only estimates a super set of the undirected edges and it only conditions on at most one node other than the pair of nodes being tested, which dramatically reduces the number of observations needed to make the test results robust. In stage two, we designed an adaptive sequential Monte Carlo (adSMC) approach to search for the optimal BN structure based on the constructed skeleton by optimizing a score function. SMC has been successfully adopted to solve optimization problems in the past (Grassberger 1997, Liu and Chen 1998, Zhang and Liu 2002, Zhang, Lin, Chen, Liang and Liu 2007, Liu 2008, Zhang, et al. 2009). Compared to other search methods, SMC is less likely to be trapped in local optima (Liu, et al. 1998, Liu 2008). Since in most SMC simulation, multiple independent instances (in our study, one instance is a fully grown BN starting from a single node) need to be generated, another advantage of SMC is that it can be run in parallel for each independent SMC instance, making it suitable for distributed or GPU-based implementations. After these two stages, we enhanced the traditional two-stage approach by adding a third stage which adds possible missed edges (not necessarily true edges) back into the network using Random Order Hill Climbing (ROHC).

Using several datasets, including simulated data, benchmark BN networks and real biological data, we show that GRASP out performed other BN structure learning methods on all these datasets. Specifically, using a heterogeneous genomic dataset containing RNA-seq, protein expressions, DNA methylations and miRNA-seq data, we applied GRASP to learn an integrated network to shed light on the function of an interesting long non-coding RNA discovered in a previous study. It demonstrated the power of GRASP in discovering novel biological relationships in integrative genomics studies.

## 2. Method

### 2.1 Preliminary

Let us denote the set of *p* variables (nodes) as ***X*** = {*X*_1_,…,*X*_*p*_}, and the set of edges as *E* = {*X*_*i*_ →*X*_*j*_} where *X*_*i*_→*X*_*j*_ are directed edges in the graph. *X*_*i*_ is called a parent of *X*_*j*_ and *X*_*j*_ a child of *X*_*i*_. Thus a graph can be represented as *G*(***X***,***E***) (Darwiche 2009).

#### Definition 1

For a set of nodes 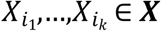 if the edges 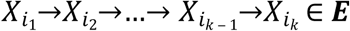, then we say 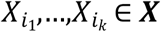 forms a directed path between 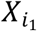 and 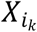.If 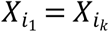, then this directed path is called a cycle.

#### Definition 2

A directed acyclic graph (DAG) is a graph *G*(*X,E*) such that all edges in *E* are directed and there is no cycles in *G*.

We denote *P*(***X***) as a joint probability distribution of the random variables in ***X***, and *Pa*_*G*_(*X*_*i*_) as the set of parents of *X*_*i*_ ∈ ***X*** given DAG *G*(***X***,***E***).

**Property 1:** *P*(***X***) can be factorized over some *G* as

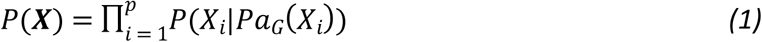

Now, we can define Bayesian network (BN) as follows:

#### Definition 3

The pair (*G,P*) is defined as a Bayesian network where the joint probability distribution *P* factorizes over *G*.

#### Remark 1

The factorization allowed the network to be locally learned, e.g. each *P*(*X*_*i*_|*Pa*_*G*_(*X*_*i*_)) can be learned independently, which saves a lot of computational time.

The factorization can usually be done in multiple different ways, that is, for some joint probability function, *P*, there exists at least two DAGs *G*_1_ and *G*_2_ that *P* factorizes over both *G*_1_ and *G*_2_.

#### Definition 4

***Q***(*P*) defines an equivalent class of *P*, containing all possible DAGs, which *P* can factorize over.

In this work, we focus on estimating any *G* ∈ ***Q*** instead of estimating every DAGs in ***Q***.

Now let us define the conditional dependencies and independencies. We denote *ind*_*P*_(*X*_*i*_;*X*_*j*_| ***Z***) as *X*_*i*_ and *X*_*j*_ are conditionally independent given ***Z*** ⊂ ***X*** with respect to *P*(***X***), and *Dep*_*P*_(*X*_*i*_;*X*_*j*_ |***Z***) as *X*_*i*_ and *X*_*j*_ are conditionally dependent given ***Z*** with respect to *P*(***X***). Here ***Z*** is a subset of variables in ***X***.

#### Definition 5

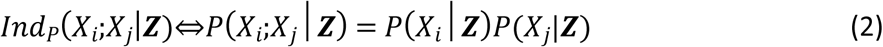

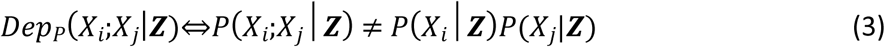

One of the most important assumptions we need to include is the faithfulness (Darwiche 2009). To define the faithfulness, let us first define trail:

#### Definition 6

A set of nodes 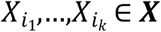 forms a trail in the graph *G*(***X***,***E***) if for every 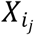 and 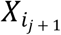, either 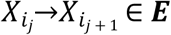 or 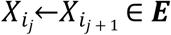.

Before we can define an active trail, let us first define descendant. If *X*_*j*_ is a descendant of *X*_*i*_ then there is a directed path from *X*_*i*_ and *X*_*j*_.

#### Definition 7

Let *G*(***X***,***E***) be a BN structure, 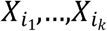 forms a trail in *G* and ***Z*** ⊂ ***X***. The trail 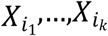 is active given ***Z*** if

- whenever there is a v-structure: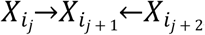, then 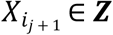 or a descendant of 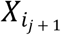 in ***Z***
- other nodes are not in ***Z***

We need one last definition, d-separation, before we can define faithfulness.

#### Definition 8

In graph *G*(***X***,***E***), for *X*_*i*_, *X*_*j*_ ∈ ***X*** and ***Z*** ⊂ ***X***, we say *X*_*i*_ and *X*_*j*_ are d-separated by ***Z***, denoted as *Dsep*_G_(*X*_*i*_;*X*_*j*_|***Z***), if none of the trails between *X*_*i*_ and *X*_*j*_ is active given ***Z***.

Now let us give the definition on the faithfulness,

#### Definition 9

*P*(***X***) is faithful to *G*(***X***,***E***) if for any ***Z*** ∈ ***X*:**

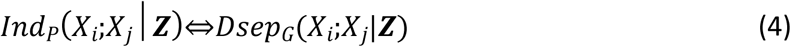

Under the faithfulness assumption, the terms conditionally independence and d-separation are equivalent; thus, they will be used interchangeably through the rest of this article.

### 2.2. The GRASP method

GRASP (GRowth-based Approach with Staged Pruning) is a three-stage algorithm for learning the structure of a BN. In the pruning stage, we designed a Double Filtering (DF) method to find the cover of the skeleton of the BN, where the skeleton of a BN is defined as the BN structure after removing the direction of all the edges, and the cover is defined as a superset of undirected edges containing all the edges of the skeleton. In the structure discovering stage, we developed an adaptive sequential Monte Carlo (adSMC) method to search the BN structure with optimal BIC score based on the undirected network found in the first stage. To reclaim the potentially missed edges, we designed a Random Ordered Hill Climbing (ROHC) method as the third stage.

#### 2.2.1. First stage: a Double Filtering (DF) method to infer the skeleton

At the beginning, for any given node, *X*, any other nodes can be its neighbors (parents or children). This first stage is to reduce the sets of neighbors for all the nodes to much smaller sets. The first filtering of the double filtering method was done by unconditioned dependency tests, filtering out the nodes that were not ancestors or descendants of a given node *X*_*i*_. The second filtering was built on dependency tests conditioning on neighboring nodes, further filtering out additional nodes as the neighbors (parents or children) of *X*_*i*_.

Let *nbr*_G_(*X*_*i*_) be the set of nodes that have an undirected edge with *X*_*i*_ (we can use *nbr*(*X*_*i*_) for convenience, if it does not cause ambiguity); the formulation of the DF method follows:

1. **First filtering:** conduct an unconditioned dependency test for each pair of nodes (*X*_*i*_ and *X*_*j*_, *i* ≠ *j*). We used mutual information test (Campos 2006) in our study, but other similar tests could be applied as well. Record the resulting *p*-value as *p*_*ij*_.If *p*_*ij*_ < *α*, update *nbr*(*X*_*i*_) as *nbr*(*X*_*i*_) ∪ {*X*_*j*_}. *α* is the predefined significance level for the test. Sort each 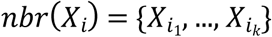, where *i*_1_, …, *i*_*k*_ ∈ {1,2, …, *p*} to obtain 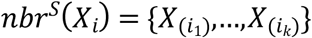, so that *p*_*i*(*i*_1_)_ ≤ *p*_*i*(*i*_2_)_ ≤,…, *p*_*i*(*i*_*k*_)_ and (*i*_1_),…,(*i*_*k*_) is a permutation of *i*_1_,…,*i*_*k*_.
2. **Second filtering**: update *nbr*^*S*^(*X*_*i*_) for each *i* ∈ {1,2,…, *p*} as follows:
  a. For node 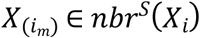 not marked as removed (starting from the one with the smallest p-value), let 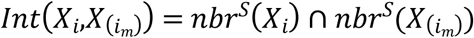. For simplicity, define 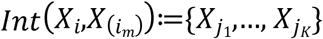, where *j*_1_,…*j*_K_ ∈ {*i*_1_,…,*i*_*k*_} \ {(*i*_m_)}.
  b. If 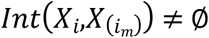, perform conditional dependency test for *X*_*i*_ and each 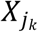 given 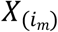 using mutual information test (Campos 2006). If the *p*-value *> α*, mark 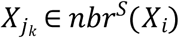 as removed.
  c. If 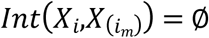, then move on to 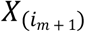 and start over from (a) until every node in *nbr*^*S*^(*X*_*i*_) is consumed.
3. **Result**: *nbr*^*C*^(*X*_*i*_) = {*X*_*j*_|*X*_*j*_ ∈ *nbr*(*X*_*i*_) and *X*_*j*_ not marked removed} is the final result of the Double Filtering. This is done for all the node in *G*.

##### Remark 2

*nbr*^*C*^(*X*_*i*_) *can be written as*

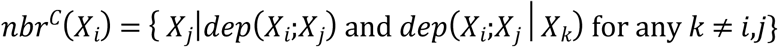

In practice, a symmetric correction may be used, where if *X*_*j*_ ∈ *nbr*^*C*^(*X*_*i*_), but *X*_*i*_ ∉ *nbr*^*C*^(*X*_*j*_), then *nbr*^*C*^(*X*_*i*_) = *nbr*^*C*^(*X*_*i*_) \ {*X*_*j*_}.

##### Theorem 1

*If the statistical power of both the marginal and conditional tests converges to 100% as the sample size n*→℞, *the resulted nbr*^*C*^(*X*_*i*_), *i* ∈ {1,2,…,*p*} *contains all true neighbors (e*.*g. true parents and children) of X*_*i*_.

The proof of this theorem is provided in Supplementary File.

#### 2.2.2. Second stage: structure searching

On the pruned space, where the sets of possible neighbors for each node have been greatly reduced, we designed an adaptive sequential Monte Carlo (adSMC) method to search the structure of the Bayesian network (*G*(***X***,***E***)). In a traditional sequential Monte Carlo, random variable X ∈ R^*d*^ is decomposed into(x_1_, x_2_, …, x_**K**_) where 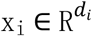 and 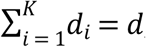, and the order of the variables to be sampled is predefined and fixed throughout the SMC sampling process. One usually samples ***x***_**1**_ first, then samples ***x***_**2**_, and so on. However, the sequence from which each variable is sampled (called sampling sequence in this study) based on any prior decomposition may not be the most efficient one. Efficiency may be gained by determining the sampling sequence dynamically. For example, when x_1_, x_2_,…, x_m_ have been sampled, where 0 < *m* < *d*, the conditional distribution *f*(x_*m* + 1_| x_1_, x_2_,…, x_m_) may have a small set of candidate decompositions (to satisfy the acyclic condition) which limits the diversity of the SMC samples. Therefore, we designed our sampling block ***x***_*i*_ conditioning on the partially sampled structure x_1_, x_2_,…, x_*i* − 1_ to increase the diversity and quality of generated samples (see **Figure S1** in Supplementary File for an example in BN).

Each SMC sample starts with all possible fully and partially connected triplets (three nodes connected by three and two undirected edges, respectively) discovered earlier in the edge screening stage, by sampling one such triplet having the least outside connection, e.g. the one having least undirected edges connected to its nodes (**Figure 1A**). These triplets are likely to be restricted to certain configuration by the sampled structure; therefore, sampling them in the early stage allows more diversity in their configurations.

**Figure 1:**
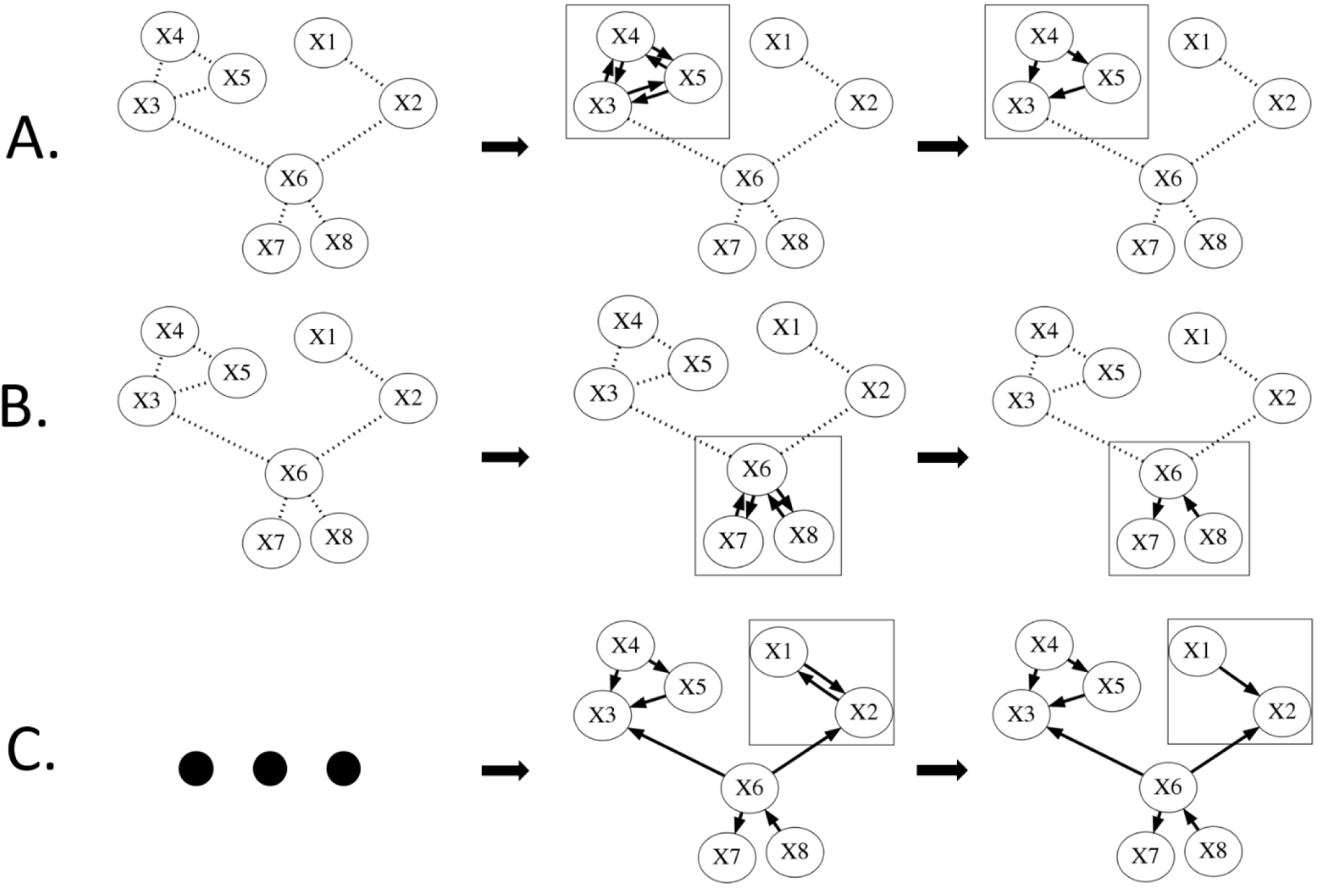
Bayesian network structure sampling procedure. (A). Each SMC sample starts with all possible fully and partially connected triplets (three nodes (i.e. X3, X4 and X5) connected by three and two undirected edges, respectively) discovered earlier in the edge screening stage, by sampling one such triplet having the least outside connection, e.g. the one having least undirected edges connected to its nodes. (B). When all the fully connected triplets were sampled, the partially connected triplets (two undirected edges among three nodes) were considered. (C). At last, we considered the pairs (the remaining undirected edges). For partially connected triplets and pairs, the configurations of the least outside connected ones were sampled first.

When all the fully connected triplets were sampled, the partially connected triplets (two undirected edges among three nodes) were considered (**Figure 1B**). At last, we considered the pairs (the remaining undirected edges, **Figure 1C**). For partially connected triplets and pairs, the configurations of the least outside connected ones were sampled first.

The probabilities of possible configurations of triplets and pairs are proportional to their BIC (Bayesian Information Criterion) score defined as 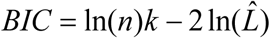, where *n* is the sample size, *k* is the number of parameters of the partial BN model, and 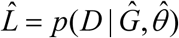 is the maximized value of the likelihood function of the BN structure, 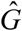, with estimated parameters, 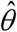, from observed data *D*. The details of the likelihood function are given in Supplementary File. The probability is set to be 0 if certain configuration fails to satisfy the acyclicity condition.

The main algorithm of this stage is summarized as follows.

1. First we consider the triplets (*X*_*I*_,*X*_*J*_,*X*_*K*_). Find the set of nodes *S* such that,

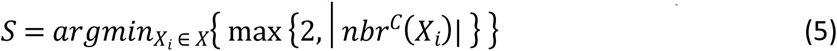

where |·| denotes the cardinality of a set. If *S* = ∅ then skip to step (6). Otherwise sample one node from *S* uniformly, say *X*_*I*_ is sampled.
2. Sample one node from 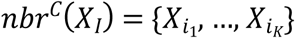 with probabilities

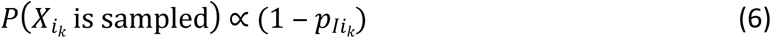 Where 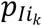 is the *p*-value from the first filtering. Suppose *X*_*J*_ is the sampled node, then remove *X*_*J*_ from *nbr*^*C*^(*X*_*I*_) and *X*_*I*_ from *nbr*^*C*^(*X*_*J*_).
3. Sample *X*_*K*_ by the following, let *INT* be the set of nodes being neighbor with both *X*_*I*_ and *X*_*J*_, *INT* = *nbr*^*C*^(*X*_*I*_) *∩ nbr*^*C*^(*X*_*J*_):
  a. if *INT* = ∅, sample *X*_K_ from *nbr*^*C*^(*X*_I_), with probability:

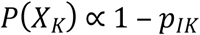
  b. if *INT* ≠ ∅, sample *X*_K_ uniformly from *INT*.
  c. remove *X*_*K*_ from *nbr*^*C*^(*X*_*I*_) and *nbr*^*C*^(*X*_*J*_) (if applicable), and *X*_*I*_,*X*_*J*_ from *nbr*^*C*^(*X*_K_).
4. Calculate the BIC for all possible configurations of the triplet {*X*_*I*_,*X*_*J*_,*X*_*K*_}. Sample one configuration with probability:

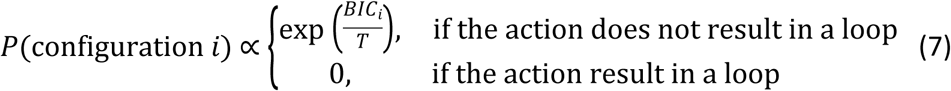 Where *T* is temperature, controlling how greedy we want the searching to be. Lower temperature will favor the configurations with larger BIC more.
5. Repeat (1)-(4) until *S* = ∅.
6. Now we handle the pairs. Find the set of node *A*, such that ∀*X*_*i*_ ∈ *A*,

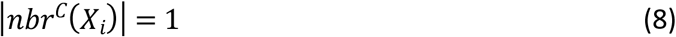

if *A* = ∅ we say this chain has converged. While *A* ≠ ∅, sample one node uniformly from *A*, say *X*_*I*_ is sampled and *X*_*J*_is the corresponding neighbor. Remove *X*_*I*_ and *X*_*J*_ from *A*.
7. Compute BIC score for any possible direction: (1). *X*_*I*_→*X*_J_, (2).*X*_I_←*X*_J_, and (3). no edge. Sample one direction with probabilities

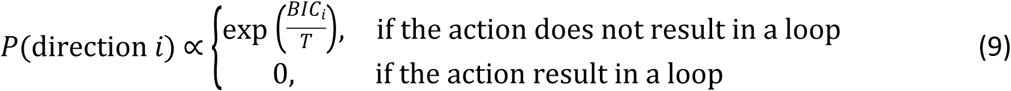

where *T* is the same as in step (4).
8. Repeat (6) and (7) until convergence (*A* = ∅). Since each SMC sample (one BN structure sampled starting from a single node) is generated independently, we can run our algorithm in parallel on multiple CPUs/GPU cores to speed up the sampling process.

#### 2.2.3. Reclaiming Missed Edges

As mentioned earlier, one disadvantage of the traditional two-stage method was that the edges missed in the first stage will never be reclaimed. Therefore, in the third stage we designed a Random Order Hill Climbing (ROHC) method to identify the possibly missed edges and refine the network. The general idea is described as following:

1. Generate a permutation of 1,2,…, *p* for each network generated by SMC, suppose *B* = *m*_1_,…, *m*_*p*_ is such a permutation.

2. For every *X*_*i*_ ∈ *X*, iterate *j* from *m*_1_ through *m*_*p*_. If *X*_*i*_←*X*_*j*_ does not create loop and results in an increasing in BIC, then we add edge *X*_*i*_←*X*_*j*_.

3. Repeat (2) until there is no possible edge to add or the searching limit is reached. One could also view this stage as a further ascent to the local optima to ensure we have the best possible BN structures.

## 3. Results

### 3.1. Benchmark networks

The networks used to generate simulated data are from actual decision making process of a wide range of real applications including risk management, tech support, and disease diagnosis (**Table 1**). All networks are obtained from Bayesian Network Repository maintained by M. Scutari http://www.bnlearn.com/bnrepository/.

**Table 1:** Bayesian networks used in the simulation study.

| **Name** | **# of nodes** | **# of edges** | **# of parameters** | **max in-degree** |
| --- | --- | --- | --- | --- |
| **Alarm** | 37 | 46 | 509 | 4 |
| **Andes** | 223 | 338 | 1157 | 6 |
| **Child** | 20 | 25 | 230 | 2 |
| **Hailfinder** | 56 | 66 | 2656 | 4 |
| **Hepar2** | 70 | 1236 | 1453 | 6 |
| **Insurance** | 27 | 52 | 984 | 3 |
| **Win95pts** | 76 | 112 | 574 | 7 |

### 3.2. Simulated data

We randomly generated data with 1000, 2000, and 5000 observations, and we generated 10 datasets for each size of observations. All results reported in this section are based on averages of the 10 datasets. Observation size in this article refers to the number of data points, and shall not be confused with number of sequential Monte Carlo samples. The datasets were generated using R package *bnlearn* using default parameters (Scutari 2009, Nagarajan, Scutari and Lèbre 2013).

### 3.3. Performance evaluation

To measure the effectiveness of edge screening methods, we employed the precision, recall and f-score measurements. Precision is defined as TP/(TP+FP), recall is defined as TP/(TP+FN), and f-score is the harmonic mean of precision and recall, 2(precision × recall)/(precision+recall), where TP means true positive (number of true undirected edges identified), FP false positive (number of non-edges identified as undirected edges), and FN false negative (number of undirected edges not identified).

In our study, recall measures the percentage of true edges (irrespective of their directions) identified; therefore, it is the most important measurement in edge screening stage, since as we discussed earlier, any missed edges in stage one may never be reclaimed in a traditional two stage approach. Besides the recall, f-score is also important since it measures a balanced performance in terms of both precision and recall. It is obvious that if we propose all possible edges, we will always identify all true edges, but that will not do any pruning to the search space. Thus, a high f-score is desired for a decent edge screening strategy.

We used Bayesian Information Criterion (BIC) as the score function in both second stage and third stage. BIC has the score-equivalent property (Darwiche 2009) (Definition 10, see below), which can reduce the search space, since if we could find one network in the equivalent class, we found the true network. And the consistency property of BIC score guarantees that the true network has the highest score asymptotically.

#### Definition 10

A score function *S*(*G*) is said to have score-equivalent property if *S*(*G*) do not distinguish among equivalent networks. That is, two Bayesian networks, G_1_ and *G*_2_, are equivalent if and only if *S*(*G*_1_) = *S*(*G*_2_).

### 3.4. Edge Screening

The principle of the edge screening stage is pruning the search space as much as possible while the remaining edges in the pruned space still possess as many true edges as possible. We compare our method to five other methods including max-min parent-child (mmpc) (Tsamardinos, et al. 2006), grow-shrink (gs) (Margaritis 2003), incremental association (iamb) (Tsamardinos, et al. 2003), fast iamb, and inter iamb (Yaramakala, et al. 2005). For all methods, we fixed the significant level (α) to 0.01.

The simulation study results (**Figure S2** and **S3** in Supplementary File) showed that our double filtering (DF) method was able to identify the most edges (highest recall) for each of the observation size we tested. In some cases we observed that with even 1000 observations, our method achieved a higher recall than the other methods using 5000 observations and the f-scores are still comparable (e.g. Alarm, Hepar2 and etc.). For some networks (Child, Insurance), not only the recalls were higher but also the f-scores were higher for DF. The results confirmed that DF identifies true edges more accurately than other methods and it often requires fewer observations. Higher recall is desired in the first stage (the edge screening stage) since any missed edges will not be recovered in the second stage.

### 3.5. Network structure sampling

There are three major factors that may affect the performance of the optimization stages (structure searching and edge reclaiming): temperature, number of SMC samples, and rounds of ROHC. Here we will discuss the effect of each of the factors, and give a general guide on how to tune these parameters.

#### 3.5.1. Temperature

The temperature parameter in SMC has the same effect as that in MCMC (Markov Chain Monte Carlo) simulations. A lower temperature will cause searching to become greedier, and higher temperatures make it less greedy. According to formula (7) and (9), when *T*→0 the searching procedure becomes a local greedy search. On the other hand, whenT→∞, the configuration is sampled uniformly. The optimal temperature is usually a value in between.

In this simulation study, we fixed SMC sample size to 20,000, and rounds of ROHC to 5. The temperature was set to between 10^−7^ and 10^−1^, increased by 10-time each time (**Figure S4** in Supplementary File). The performance is shown in the relative scale (BIC of true network/BIC of the learned network), where higher ratio means higher BIC score; thus, better network structure. Lower temperatures in most cases gave a lower score, as well as higher temperatures, consistent with what we would expect. Most of the optimal scores happened around T = 0.001 or 0.01. We can also see that the optimal temperature does not depend on the observation sizes, since the optimal temperatures are the same across the 3 different observation sizes. Another observation we had was that the optimal temperatures did not change much when the number of variables (nodes) changes. From **Figure S4** we can see that for Andes (with 223 nodes) and child (20 nodes), the optimal temperature is both around 0.01 and 0.001.

#### 3.5.2. SMC samples

In general, generating more SMC samples gives a higher chance to reach the optimum. However, more samples also require more computation time; therefore, a balance between running time and sample sizes must be made. In most of our simulation study and practical problems, we found that around 20,000 samples were often good enough for finding a network with a relatively high BIC score.

#### 3.5.3. Adaptive SMC

To show the improvement of using adaptive SMC, we compared the BICs of 20,000 SMC samples between the adSMC and traditional SMC (**Figure S5** in Supplementary File). In the traditional SMC, we designed the sampling block in the order of fully connected triplets, partially connected triplets and pairs, and started from least outside connected ones. Clearly, the adSMC generates higher scored networks in general.

### 3.6. Necessity of reclaiming

We discussed earlier that there could be some true edges missed in the first stage due to the test power and data quality. Here we will show that Random Order Hill Climbing (ROHC) indeed improves the learned BN structure. We used *alarm* and *win95pts* networks to illustrate the improvement made by ROHC (**Figure S6** in Supplementary File). They both had significance level cut off of 0.01, temperature 0.001, and 20,000 SMC samples. As we can see, the improvements were substantial. The optimum network found by SMC was also improved. Therefore, it is necessary to have the third stage to further refine the learned network.

However, one should notice that the complexity level of ROHC is approximately *O*(*N*^2^); therefore, in a typical network with hundreds of nodes only 1 or 2 rounds of ROHC are affordable.

### 3.7. Overall performance

We evaluated the overall performance of our method and the general two stage methods (5 edge screening methods, gs, mmpc, iamb, fast.iamb, and inter.iamb combined with 2 optimization methods, Hill climbing and tabu search) on 7 benchmark networks. The results are shown in **Figure 2**. For 3 different observation sizes, our method out performed all the general two-stage methods on almost all benchmark networks except on the hepar2 network where all methods achieved similar scores, which are very close to the BIC of the true network.

**Figure 2:**
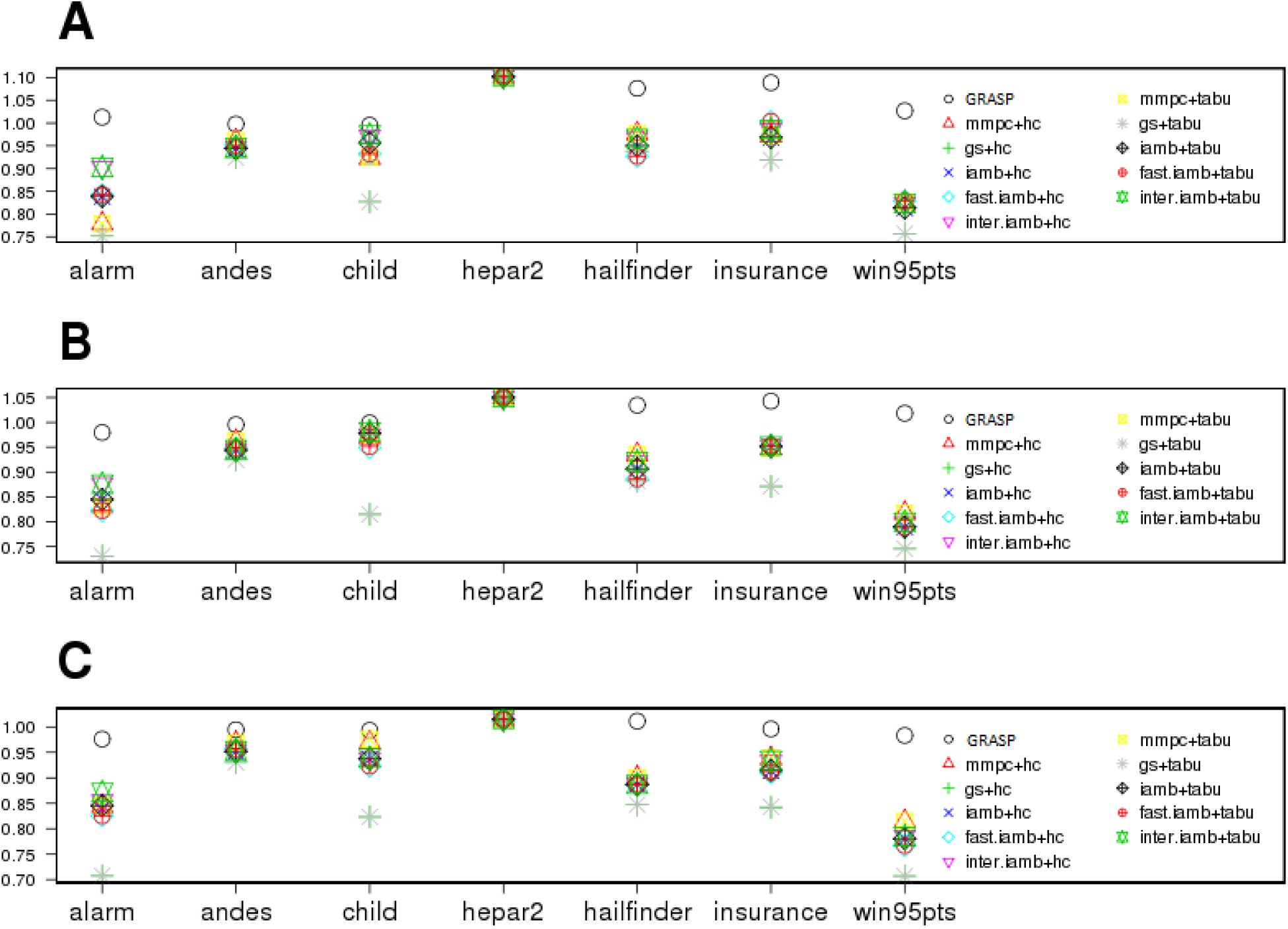
BIC scores of all methods on 7 benchmark networks. Observation sizes: (A) 1000, (B) 2000, (C) 5000. GRASP achieved highest BIC scores on most of the network except hepar2, for which all methods performed well.

### 3.8. Real data study

#### Flow cytometry dataset

We applied our method to the flow cytometry dataset (Sachs, Perez, Pe’er, Lauffenburger and Nolan 2005). There are 11 proteins and phospholipid components of the signaling network. The original data were collected from 7466 cells, containing continuous values for the 11 components. Sachs et al suggested to get rid of the potential outliers by removing data that are 3 standard deviations away from any attribute. Thus the data we are analyzing contains 6814 observations, each with 11 values corresponding to 11 variables (nodes in the network). We discretized each variable into 3 categories, for high/medium/low levels, with each level containing 33% of the data. We compared our method to the general 2-stage methods and the CD method (Fu, et al. 2013). We can see from **Figure 3** that our method has the highest BIC score, consistent with our observation in simulation study that our method could fit the data with better models.

**Figure 3:**
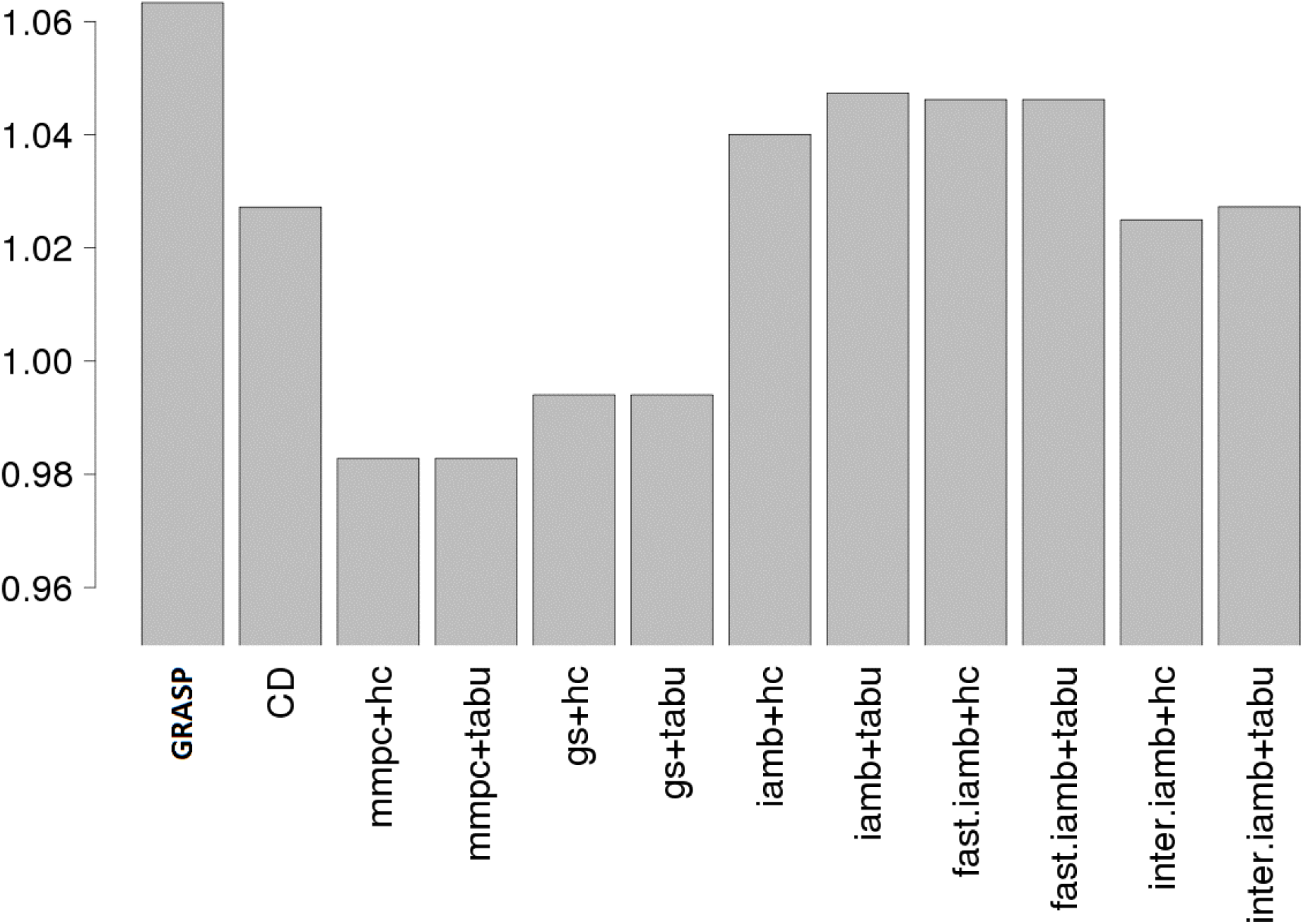
BIC scores for the flow cytometry data from 12 methods. GRASP has the highest BIC score.

#### An integrative genomic study using the cancer genome atlas (TCGA) data

In a previous study of ours (Stewart, Luks, Roycik, Sang and Zhang 2013), we have identified a long non-coding RNA, LOC90784, which is strongly associated with breast cancer health disparity between African American and Caucasian American breast cancer patients. However, literature search resulted in no information about it since it had not been studied by any researchers in the past. Using several different types of genomics data, we applied GRASP to perform an integrative study to build a Bayesian network with different genomics features to shed some light on the function of this transcript. We first used RNA-seq data to identify transcripts highly correlated with LOC90784. There are thousands of genes with significant correlation coefficients with LOC90784 as measured by adjusted p-values (padj <= 0.05). We chose a relatively large correlation coefficient cutoff to give us a small number of genes to control the size of resulting networks. Eight transcripts were selected with absolute values of correlation coefficients greater than 0.27. We then found other genomic features, including microRNAs, DNA methylations and protein expressions that are highly correlated with these transcripts, which gave us 13 microRNAs, 5 DNA methylation regions (aggregated around genes) and 5 proteins. Using the samples with all the above measurements, we inferred the BN structure for these genomics features as shown in **Figure 4**. Before applying GRASP, continuous variables were discretized into 2 categories by their medians. As a comparison, bnlearn produced a network without LOC90784 (**Figure S7** in Supplementary File). **Figure 4** shows rather complex relationships among all these genomic features. A thorough investigation of this network is beyond the scope of this work. However, some literature search on the nodes around LOC90784 provided interesting hypotheses, which could be followed up with experiments. Specifically, TET3, an upstream gene, was found to inhibit TGF-β1-induced epithelial-mesenchymal transition in ovarian cancer cells (Ye, et al. 2016). High frequency of PIK3R2 mutations in endometrial cancer was found to be related to the regulation of protein stability of PTEN (Cheung, et al. 2011), which is a well-known cancer related gene. There are not a lot of published studies on IRGQ. From the Human Protein Atlas database (https://www.proteinatlas.org/ENSG00000167378-IRGQ/pathology) we found that this gene is a prognostic biomarker and significant for survival for several cancer types including pancreatic cancer, renal cancer, cervical cancer and liver cancer. It would be interesting to see how perturbations of TET3, PIK3R2, such as knockdown/knockout experiments, affect LOC90784 and how perturbation of LOC90784 affects IRGQ. If we trace upstream of these genes, we will find other genomics features such as DNA methylations or miRNAs that regulate either TET3 or PIK3R2. These hypotheses demonstrated the potential of GRASP for discovering new biology through integrative genomic studies.

**Figure 4.**
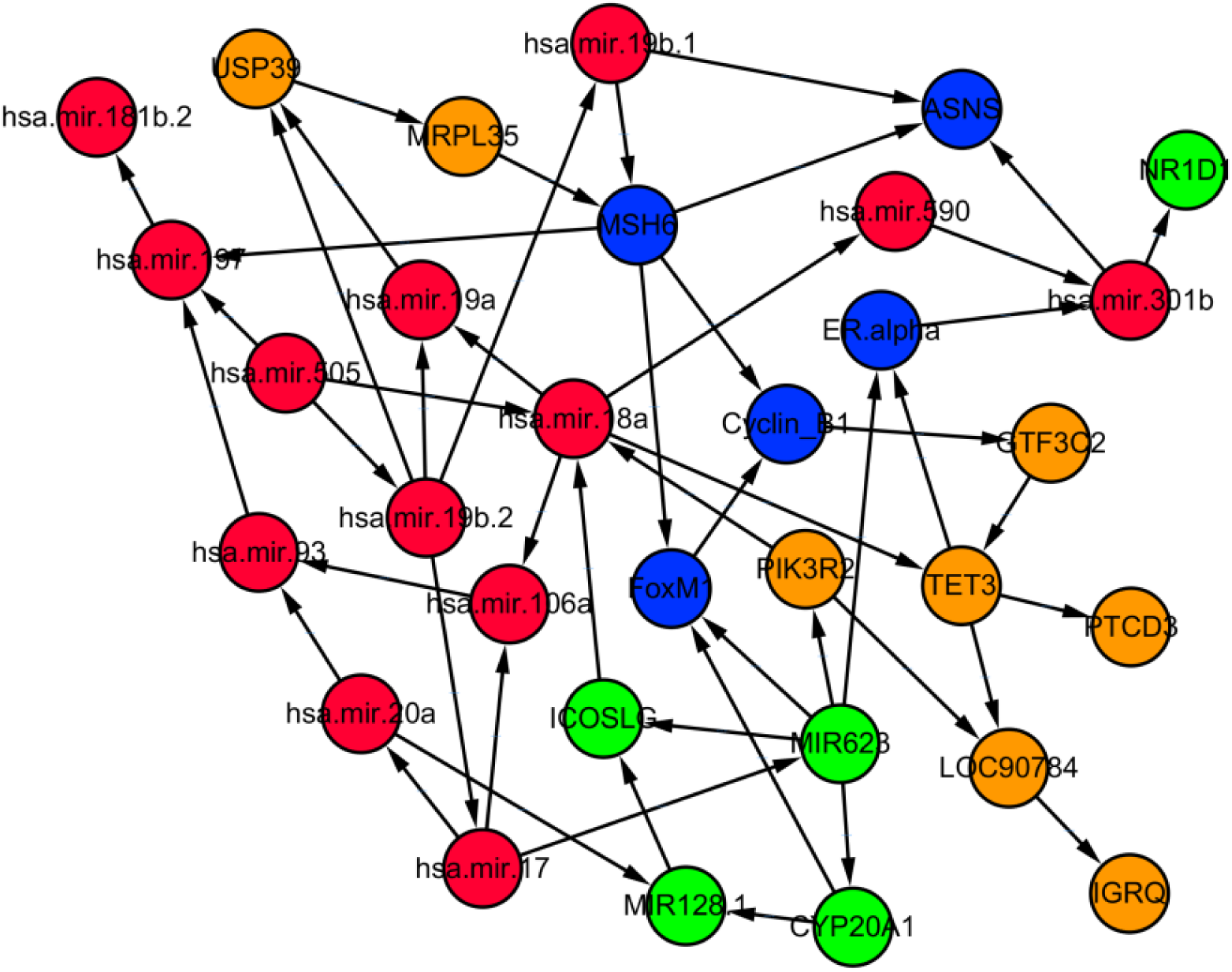
The BN structure learned using multiple different genomic features which are highly correlated with the expression of LOC90784. Orange nodes: mRNA transcripts; Red nodes: microRNAs; Blue nodes: protein expressions; Green nodes: DNA methylations.

## 4. Conclusion and Discussion

In this work, we developed a three-stage Bayesian network structure learning method, GRASP. The first stage is a new edge screening method, Double Filtering (DF), which recovers a super set of true edges with satisfactory recall and f-score. The second stage is an adaptive SMC approach to sample BN structures that optimize the score function (BIC in this study). To reclaim the possibly missed edges from the first two stages, we developed a random order hill climbing (ROHC) method to add additional edges to the BN structure sampled at the second stage to further improve the BIC score. The principle of DF is quite different from the well-known mmpc method or other similar constraint based methods, which aim to identify the exact skeleton of the BN (undirected true edges). DF focuses on identifying a set of undirected edges that contains all the true edges, while minimizing the number of false positives. The advantage of mmpc is that given enough observations it identifies the true network skeleton; however, it may not be feasible when the numbers of observations are relatively small since mmpc conducts conditional dependency test conditioning on all previously identified dependent (connected) nodes, and it requires substantially more observations when the number of conditioned nodes increases. On the other hand, DF only conditions on one node at a time, so the required observation size can be much smaller.

The adSMC approach in structure sampling stage provided us better chance to find global optimal structure than greedy search and other heuristic sampling algorithms. In addition, adSMC sampling is completely parallelizable, and multiple CPUs/GPU implementations will likely further improve the computational efficiency substantially.

Although in this study we focused on categorical variables (nodes) with multinomial distribution, one may extend our approach to other types of variables including Gaussian ones, as long as all the nodes have the same distribution and the local conditional distribution can be estimated. Imposing distributions that are easier to be estimated on the nodes will in general make the search more efficient.

For mixed type of BNs (where nodes do not necessarily have the same distribution), our method could handle them indirectly by discretizing the observations and making each node having multinomial distributions. Learning the structures of these BNs will be an interesting research topic in a follow-up study.

## Acknowledgement

This work was partially supported by the National Institute of General Medical Sciences of the National Institute of Health under award number R01GM126558 (JZ), and the University of Rochester CTSA award number UL1 TR002001 from the National Center for Advancing Translational Sciences of the National Institutes of Health (XQ). The funders had no role in the study design, data collection and analysis, decision to publish, or preparation of the manuscript.

